# Fentanyl binds to the μ-opioid receptor via the lipid membrane and transmembrane helices

**DOI:** 10.1101/2021.02.04.429703

**Authors:** Katy J Sutcliffe, Robin A Corey, Steven J Charlton, Richard B Sessions, Graeme Henderson, Eamonn Kelly

## Abstract

Overdose deaths from synthetic opioids, such as fentanyl, have reached epidemic proportions in the USA and are increasing worldwide. Fentanyl is a potent opioid agonist, that is less well reversed by naloxone than morphine. Due to fentanyl’s high lipophilicity and elongated structure we hypothesised that its unusual pharmacology may be explained by a novel binding mode to the μ-opioid receptor (MOPr).

By employing coarse-grained molecular dynamics simulations and free energy calculations, we determined the routes by which fentanyl and morphine access the orthosteric pocket of MOPr.

Morphine accesses MOPr via the aqueous pathway; first binding to an extracellular vestibule, then diffusing into the orthosteric pocket. In contrast, fentanyl takes a novel route; first partitioning into the membrane, before accessing the orthosteric site by diffusing through a ligand-induced gap between the transmembrane helices.

This novel lipophilic route may explain the high potency and lower susceptibility of fentanyl to reversal by naloxone.

## Introduction

The synthetic opioid agonist, fentanyl, has been in medicinal use for over 50 years as a powerful, fast-acting analgesic and for induction of sedation and general anaesthesia. However, since 2014 fentanyl and fentanyl analogues (fentanyls) have increasingly appeared in the illicit drug market in North America (1, 2); this has been associated with a dramatic rise in the number of acute opioid overdose deaths involving fentanyls (3), with the ease of synthesis and transport making fentanyls attractive to suppliers of illicit opioids (4). Concerningly, there are increasing reports that fentanyl overdose requires higher doses of the antagonist naloxone to reverse, compared to heroin (4–10). Indeed, we have recently shown that naloxone reverses respiratory depression induced by fentanyl in mice less readily, than that induced by morphine (11). This finding is at odds with classical receptor theory, as under competitive conditions the degree of antagonism depends only on the affinity and concentration of the antagonist, not the potency of the agonist (12). Fentanyls, therefore, are an increasing public health concern, and exhibit a unique pharmacology which is yet to be fully understood.

In *in vitro* radioligand binding and signaling experiments there is a discrepancy between the relative potencies of fentanyl and morphine in experiments performed in membrane homogenate and intact cell systems. In membrane homogenate experiments fentanyl and morphine exhibit similar affinity of binding to the m opioid receptor (MOPr), both in the absence and presence of Na^+^ ions (13–15), whilst in membrane homogenate studies of receptor activation using GTPγS binding the potency of fentanyl has been reported to be less than 2 fold greater than that of morphine with fentanyl having equal or only slightly greater agonist efficacy (14–16). In marked contrast, in intact cell studies of MOPr signaling (cyclic AMP inhibition, G protein activation and arrestin translocation) fentanyl is some 5 to 50 fold more potent than morphine (16–20). We propose that this discrepancy may be explained by the unusual properties of fentanyl. Firstly, fentanyls are highly lipophilic compared to other opioid ligands (21, 22) and may therefore in intact cells interact with, or even partition into, the lipid bilayer thus increasing the concentration of drug in the vicinity of the receptor (23–25). Secondly, fentanyls have an elongated chemical structure with a central protonatable nitrogen and 6 rotatable bonds, compared to the rigid ring structure of morphine, naloxone and other morphinan compounds (Supplementary Fig 1A). This flexible structure may facilitate a novel binding process, distinct from that of morphinan opioid agonists and the antagonist naloxone.

MOPr, which mediates the pharmacological effects of fentanyl (11), is a G protein-coupled receptor (GPCR) found primarily at the plasma membrane. The MOPr structure follows the general architecture of a class A GPCR (26–28), with a deep aqueous binding pocket for orthosteric ligands which is shielded from the extracellular milieu by three extracellular loops (ECLs) and from the lipid bilayer by the seven transmembrane helices (TMDs). X-ray crystal structures and cryo-electron microscopy structures of the MOPr reveal small molecule morphinan ligands (26, 27) and the peptide DAMGO (28) bind within this orthosteric site via a key interaction between the protonated amine of the ligand and D147^3.32^ (residues are labelled according to the Ballesteros-Weinstein system (29)) (Supplementary Fig 2).

It is generally assumed that GPCR ligands bind to the orthosteric site directly from the extracellular aqueous phase (30–33). However, studies of sphingosine-1-phosphate, cannabinoid (CB2), protease activated (PAR1), and purinergic (P2Y1) receptors have shown that some highly lipophilic ligands are able to access the orthosteric pocket by diffusing through the lipid membrane and the receptor transmembrane helices (34–37). Therefore, based on fentanyl’s high lipophilicity, elongated structure, and differing potencies dependent on the membrane environment, we hypothesised that fentanyl may bind to the MOPr in a non-canonical fashion via the lipid bilayer.

Long timescale all-atom molecular dynamics (MD) simulations have been used to capture small molecule ligands binding to GPCRs (30, 32, 33). However, capturing a rare event such as ligand binding usually requires millisecond timescale simulations using specially designed machines (38). In addition, fentanyl has many rotatable bonds and therefore many degrees of freedom which can pose sampling problems. Coarse-grain (CG) MD can be utilised to overcome these sampling issues (39–41). In CG MD, rather than representing each individual atom as a defined bead, groups of atoms are represented as a single bead which describes the overall properties of the chemical group. This lower resolution representation results in fewer beads and fewer degrees of freedom to describe a system, meaning the conformational landscape can be sampled more efficiently, and rare events such as ligand binding can be more readily captured (42, 43). To determine how fentanyls and morphinans access and bind within the orthosteric site, we employed unbiased CG MD simulations of membrane-embedded MOPr solvated in water, ions and opioid ligands.

## Results

We built molecular systems of the MOPr (26, 44, 45) (PDB: 4DKL) in a solvated membrane using the coarse grained MARTINI 2.2 force field (see Methods for details) (Supplementary Fig 1B). We added 6 molecules of either protonated fentanyl, neutral fentanyl, protonated morphine or neutral morphine (Supplementary Fig 1C and D) to the solvent and ran 3 - 6 independent repeats of 1 - 5 μs, unbiased CG MD simulations to allow the ligands to bind to the MOPr (Supplementary Table 1).

We first characterised how the protonated and neutral forms of fentanyl and morphine interacted with the phospholipid membrane. In all simulations, fentanyl and morphine rapidly diffused from the solvent to interact with the phospholipid bilayer. Both the protonated and neutral fentanyl molecules fully partitioned into the membrane (Fig 1A), with the neutral form of the ligand penetrating deeper into the bilayer centre (Fig 1C and Supplementary Fig 3A). In contrast morphine interacted only with the phosphate head groups at the lipid-solvent interface (Fig 1B), and neither the protonated nor neutral form of the ligand partitioned into the bilayer (Fig 1C).

**Figure. 1.**
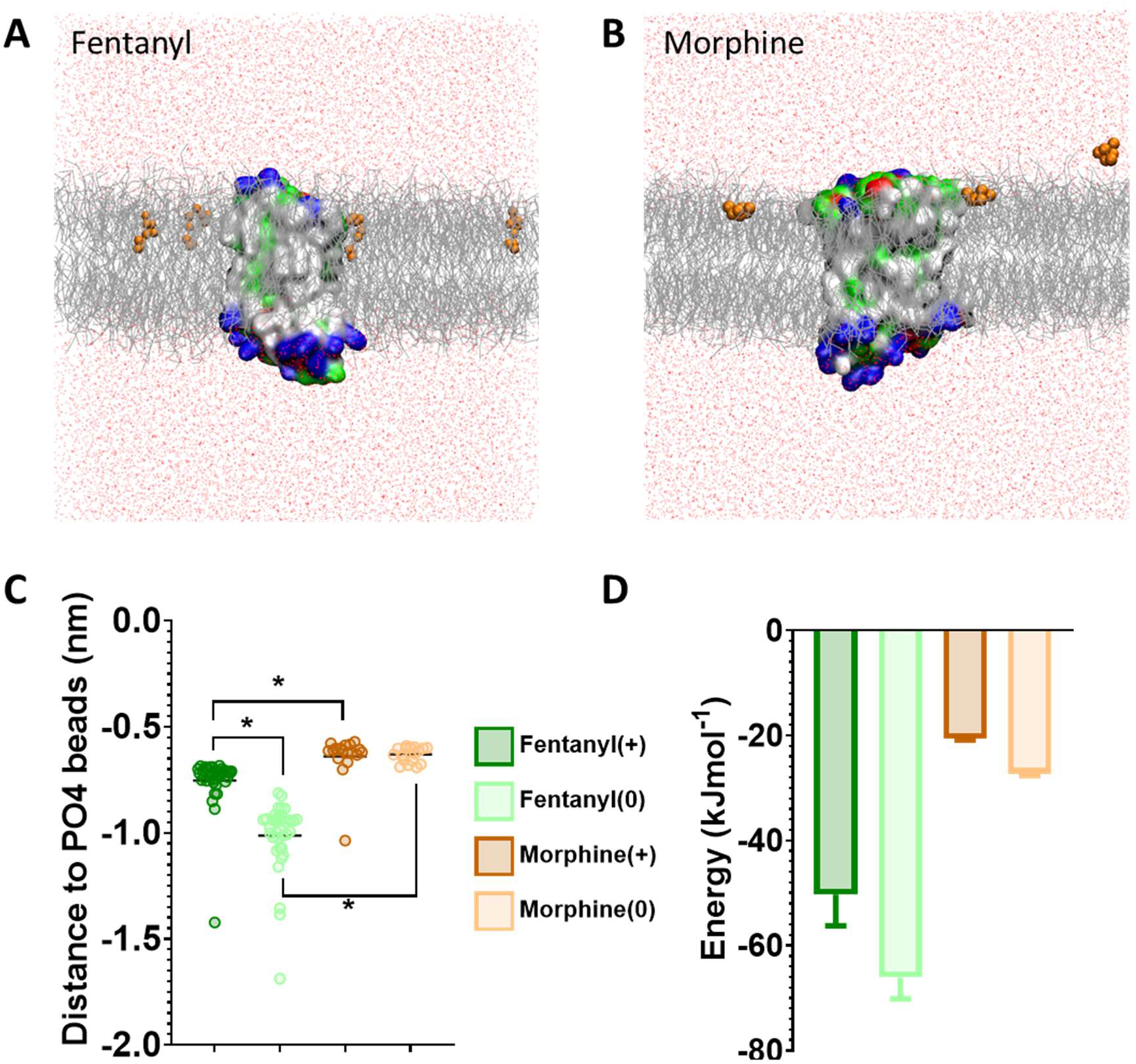
Differences in how opioid ligands partition into the lipid bilayer. **A.** Fentanyl molecules (orange) rapidly partition into the lipid membrane (grey). **B.** Morphine molecules (orange) do not fully enter the lipid membrane (grey) but interact with the charged lipid headgroups. Note while ligands can appear on either side of the bilayer due to the periodic boundary conditions applied in these simulations, for clarity only ligands in the upper leaflet of the membrane are shown. In no simulation did a ligand travel all the way through the bilayer. The protein is coloured according to residue properties (hydrophobic; grey, polar; green, acidic; red, basic; blue). **C.** Distance between the center of mass of the ligand and the phosphate head groups (PO4 beads) of the lipid bilayer. Both the charged and neutral forms of fentanyl partition significantly deeper in the membrane than morphine. * p < 0.05, one-way ANOVA. Each data point represents the average distance between a fentanyl molecule and the PO4 beads over the entire simulation. **D.** Free energy change for ligands moving between the bilayer center and the aqueous solvent. Calculated from PMF profiles shown in Supplementary Fig 3B-E. Data plotted as mean ± error calculated from bootstrap analysis.

To further quantify the propensity for fentanyl and morphine to partition between the aqueous and lipid phase, we performed steered MD and umbrella sampling to calculate the free energy change (ΔG) of membrane partitioning. Steered MD uses an external force to “pull” the ligand away from the center of the membrane (46), creating a trajectory of the ligand moving between the lipid and aqueous solvent from which umbrella sampling can be performed to extract potential of mean force (PMF) profiles. Using these PMFs, ΔG can be calculated as the free energy difference between the ligand residing in the bilayer center verses the aqueous solvent. The resulting ΔG values are shown in Fig 1D, and the PMF profiles in Supplementary Fig 3B-E.

The calculated ΔG for membrane partitioning for the protonated and neutral forms of fentanyl were −50.3 ± 6.0 kJmol^-1^ and −66.1 ± 4.1 kJmol^-1^, respectively. Whereas, the values for morphine showed a much smaller free energy difference (protonated; −20.6 ± 0.3 kJmol^-1^, neutral; −27.3 ± 0.3 kJmol^-1^). The spontaneous membrane partitioning exhibited by fentanyl in the unbiased CG simulations, along with this greater free energy change in partitioning between the lipid and the aqueous solvent, suggests that fentanyl has a greater propensity to concentrate in the cell membrane, compared to morphine.

### Fentanyl binds to MOPr via the lipid phase and the transmembrane helices

For the remaining analyses, we focus on the simulations of the protonated ligands, as the charged species is required to form the canonical amine – D147^3.32^ salt bridge essential for opioid ligand binding within the orthosteric pocket.

In the CG MD simulations fentanyl molecules in the lipid bilayer appeared to congregate around MOPr (Fig 1A). We therefore constructed ligand density maps across all the fentanyl simulations (Fig 2A), using the VMD VolMap tool (47). Fentanyl molecules cluster around the receptor helices in the upper leaflet of the membrane, with densities determined on the lipid-facing sides of the TM1/2, TM6/7 and TM7/1 interfaces.

**Figure. 2.**
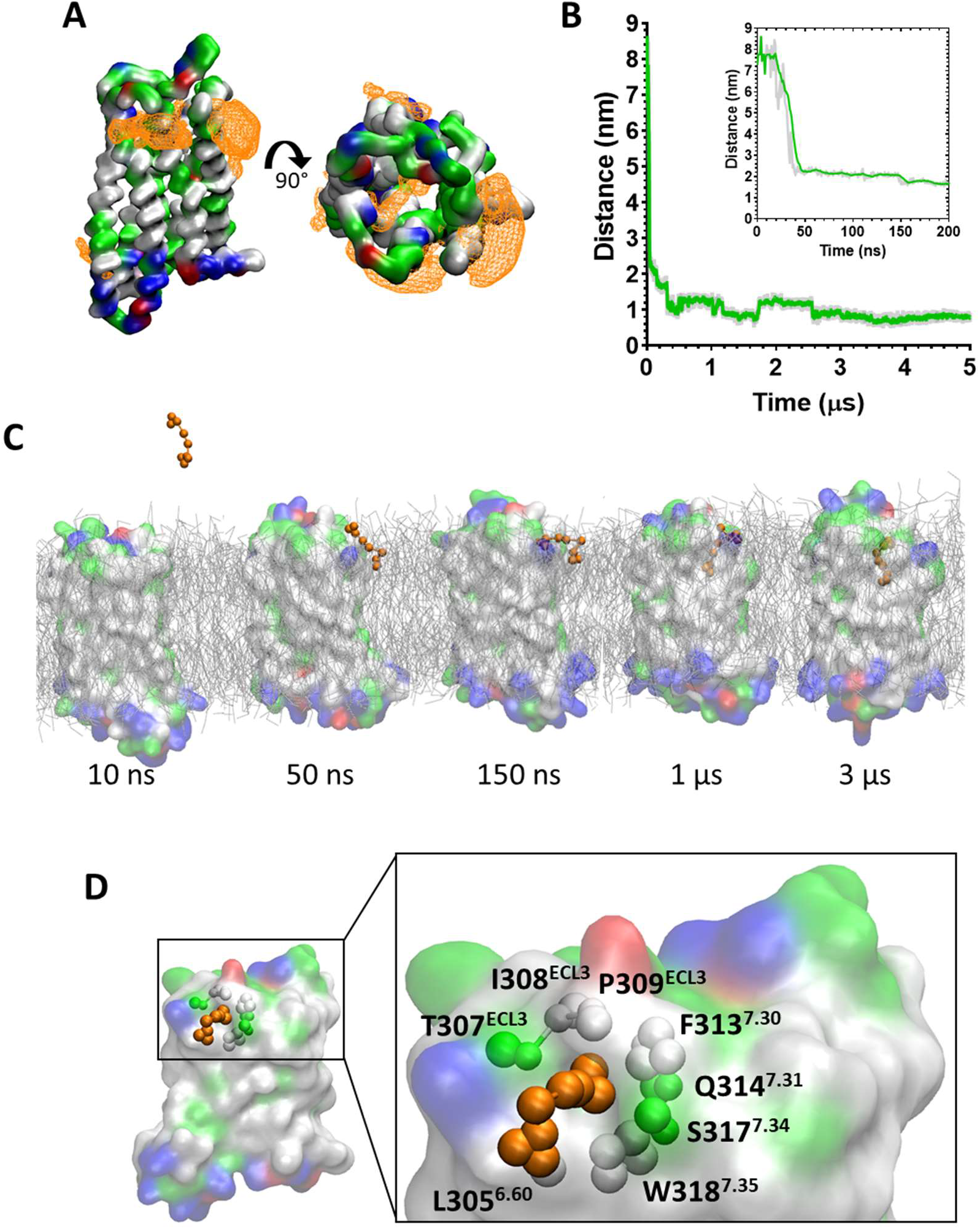
Fentanyl binds to the MOPr from the lipid phase, via a gap between TM6 and TM7. **A.** Ligand density maps averaged over the 5 μs simulation, show fentanyl densities around the receptor transmembrane domains and within the orthosteric pocket (orange). The protein is coloured according to residue properties (hydrophobic; grey, polar; green, acidic; red, basic; blue). **B.** Distance between the Qd bead of fentanyl and the SC1 bead of D147^3.32^ over the entire 5 μs and in the first 200 ns (inset). Data are presented as the raw values (grey) and moving average over 10 frames (green). **C.** Snapshots from the unbiased simulation of fentanyl binding to MOPr. Fentanyl moves from the aqueous solvent into the lipid bilayer, then interacts with the MOPr transmembrane domains and induces the formation of a gap between TM6 and 7, through which fentanyl accesses the orthosteric site. **D.** Fentanyl at the TM6/7 interface. Fentanyl is depicted as orange beads, and the residues comprising the lipid entry gap as coloured beads.

Most notably, we also observed fentanyl diffusing through MOPr to the orthosteric binding pocket via a novel lipophilic pathway (see Supplementary Movie 1). Snapshots from the MD simulation (Fig 2C and Supplementary Fig 4A) show fentanyl first partitioning into the lipid bilayer, then interacting with a ligand-induced gap (Supplementary Fig 4C and D) at the TM6/7 interface, and finally accessing the orthosteric site by diffusing through this gap in the MOPr helices. The fentanyl molecule took 3 μs to diffuse across the receptor TM domains to the orthosteric site (Fig 2B).

The TM6/7 interface and the gap induced by the fentanyl molecule is shown in Fig 2D. This interface comprises hydrophobic and polar residues from TM6 and 7, as well as ECL3. Specifically, the relatively small side chains of L305^6.60^, T307^ECL3^, I308^ECL3^ and P309^ECL3^ allow formation of a pore through which the phenethyl group of fentanyl (represented by the F1, F2 and F3 beads, see Supplementary Fig 1D) can access the receptor orthosteric pocket. Meanwhile, the aromatic side chain of W318^7.35^ stabilises the position of fentanyl’s N-phenylpropanamide (represented by the F7, F8 and F9 beads, see Supplementary Fig 1D).

### Morphine binds to MOPr via the aqueous phase and an extracellular vestibule site

During the unbiased CG simulations, we observed morphine spontaneously binding to the MOPr via the canonical aqueous pathway (see Supplementary Movie 2). Ligand density maps show a density for a morphine molecule in the extracellular portion of the MOPr; above and within the orthosteric binding site (Fig 3A). Plotting the distance between the charged Qd bead of morphine and the side chain bead of D147^3.32^ shows that the ligand rapidly diffuses from the aqueous solvent to interact with the extracellular surface of MOPr within the first 50 ns of the CG simulation (Fig 3B). Morphine maintains stable interactions with this extracellular site for 4.2 μs, before finally moving deeper into the orthosteric binding pocket. Figure 3C and Supplementary Fig 4B show snapshots of morphine travelling along this canonical aqueous binding pathway, with it initially binding to the extracellular vestibule site and then finally binding within the orthosteric pocket.

**Figure. 3.**
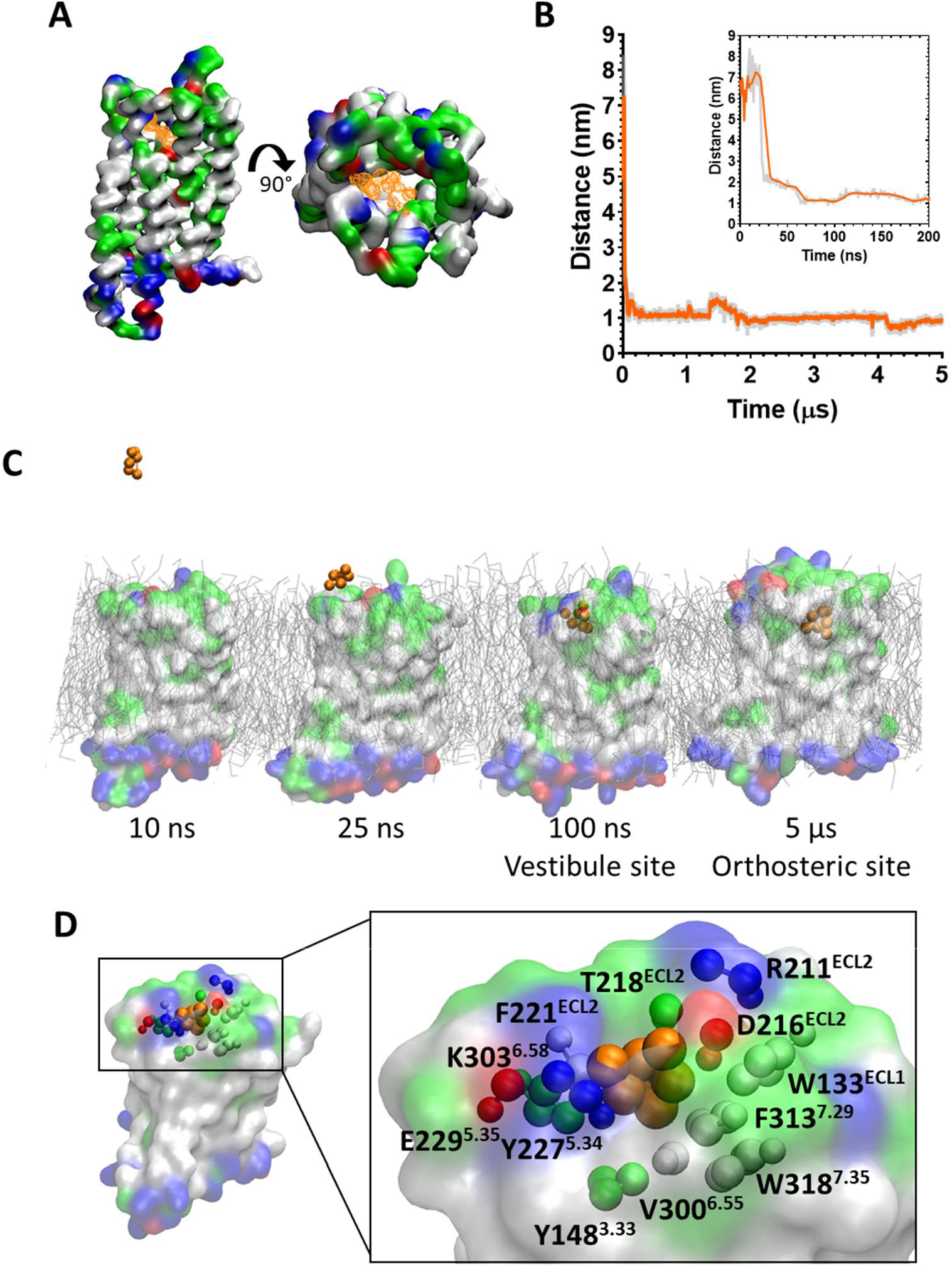
Morphine binds to the MOPr from the aqueous phase, via an extracellular vestibule site. **A.** Ligand density maps averaged over the 5 μs simulation, show morphine densities above and within the orthosteric pocket (orange). The protein is coloured according to residue properties (hydrophobic; grey, polar; green, acidic; red, basic; blue). **B.** Distance between the Qd bead of morphine and the SC1 bead of D147^3.32^ over the entire 5 μs and in the first 200 ns (inset). Data are presented as the raw values (grey) and moving average over 10 frames (orange). **C.** Snapshots from the unbiased simulation of morphine binding to MOPr. Morphine moves from the aqueous solvent to an extracellular vestibule and finally the orthosteric site. **D.** Morphine in the extracellular vestibule site. Morphine is depicted as orange beads, and the residues comprising the vestibule site as coloured beads.

The extracellular vestibule site is shown in Fig 3D, comprising primarily polar or charged residue side chains in ECL2 and the extracellular ends of TMs 5, 6 and 7. This extracellular vestibule site (48) appears to be a conserved feature of small molecule binding to Class A GPCRs, having previously been highlighted in MD simulations of the β1 and β2 adrenergic receptors (32), M3 muscarinic receptor (33), adenosine A2A receptor (42) and oliceridine binding to the MOPr (30).

### Calculation of the relative binding energies in the aqueous and lipophilic access routes

Next, we sought to further characterize the aqueous and lipid binding pathways by calculation of the free energy of binding (Δ_Gbinding_) for each ligand in each pathway.

Starting from the final frames of the simulations where fentanyl (Fig 2C) or morphine (Fig 3C) bound in the orthosteric site, steered MD simulations were performed to recreate the aqueous and lipid binding routes for each ligand. Ligands were “pulled” from the orthosteric site along either the aqueous or lipid access route, generating a trajectory from which starting conformations for umbrella sampling could be generated. The resulting PMF profiles are presented in Fig 4, along with the calculated ΔG_binding_ values for each ligand in each binding pathway. Here, ΔG_binding_ represents the free energy difference between the ligand-bound MOPr and the unbound ligand residing in either the aqueous solvent (Fig 4A and B) or the lipid membrane (Fig 4C and D).

**Figure. 4.**
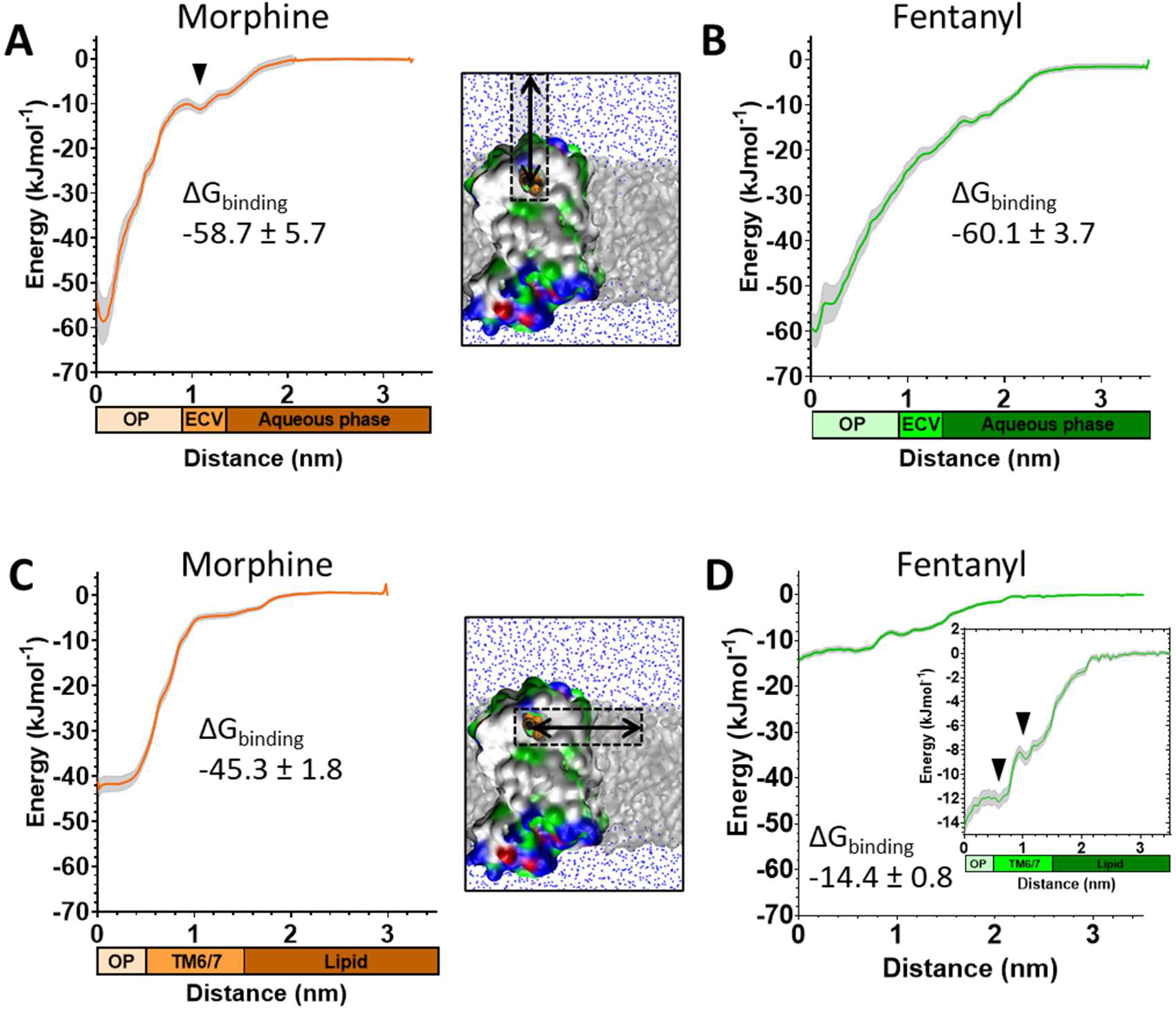
Free energy calculations for ligand binding pathways. Steered MD was used to recreate the spontaneous binding events reported in Figs 2 and 3. Umbrella sampling and the weighted histogram analysis method were then employed to determine the free energy of binding for each ligand in each pathway. In all plots the distance along the reaction coordinate is defined as the distance between the centre of mass of the ligand and receptor. Coloured bars beneath the x-axes indicate the orthosteric pocket (OP), extracellular vestibule (ECV), TM6/7 interface, lipid and aqueous phases. Data are plotted as an average (coloured line) and statistical error (grey), calculated from bootstrap analysis. δG_binding_ is expressed as mean ± statistical error. **A.** PMF profile for morphine binding via the aqueous pathway. **B.** PMF profile for fentanyl binding via the aqueous pathway. **C.** PMF profile for morphine binding via the lipid pathway. **D.** PMF profile for fentanyl binding via the lipid pathway. Inset shows the same data with expanded y axis.

The PMF profiles for morphine and fentanyl binding via the aqueous pathway are shown in Fig 4A and B, respectively. The calculated ΔG_binding_ for each ligand is similar (−58.7 ± 5.7 kJmol^-1^ for morphine, −60.1 ± 3.7 kJmol^-1^ for fentanyl), suggesting that both ligands can bind via this aqueous route with similar ease. In the profile for morphine binding a small local minimum can be seen between 1.0 - 1.3 nm, indicating the extracellular vestibule site identified in the unbiased MD simulations (Fig 2D). In the profile for fentanyl binding no small local minimum indicative of binding to the extracellular vestibule is apparent.

The PMF profiles for morphine and fentanyl binding via the lipid pathway are shown in Fig 4C and D. For morphine, the PMF profile follows a steep curve, with a calculated ΔGbinding of −45.3 ± 1.8 kJmol^-1^. In contrast, the fentanyl ΔG_binding_ is significantly lower (−14.4 ± 0.8 kJmol^-1^), with two local minima at 0 – 0.8 nm and 1.1 – 1.5 nm, corresponding to the orthosteric site and the TM6/7 interface (Fig 2D) on the lipid-facing side of the helices, respectively.

### Comparison of free energy landscapes for morphine and fentanyl

In order to compare the full binding pathways from solvent to MOPr for fentanyl and morphine, we used the data from the PMF analyses in Fig 1D and 4 to construct free energy landscapes for both ligands in their protonated forms as they interact with MOPr (Fig 5). Figure 5A shows a thermodynamic cycle for each ligand, where ΔG_1_ is free energy of transfer between the receptor and the membrane, as measured in Fig 4 C and D, ΔG_2_ is between the membrane and solvent, as per Fig 1D, ΔG_3_ is the energy of moving in the solvent (assumed to be 0 kJ mol^-1^) and ΔGdirect represents the aqueous pathway from solvent to orthosteric binding site in the receptor explored in Fig 4A and B. From this, we can state that:

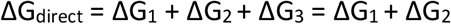

**Figure. 5.**
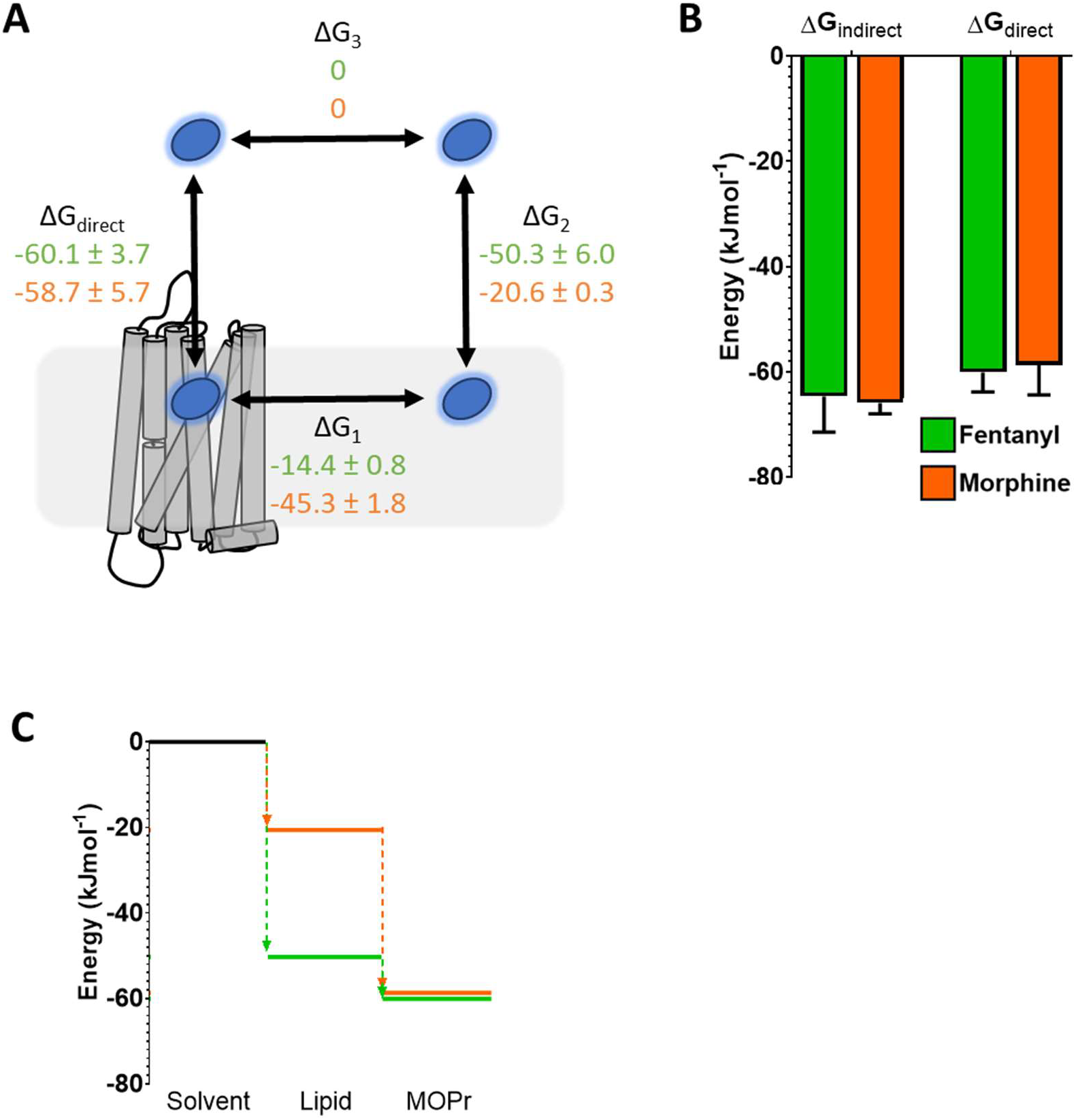
Comparison of free energy landscapes for fentanyl and morphine binding to the MOPr. **A.** Thermodynamic cycle for opioid ligand binding to MOPr; either by the direct, aqueous pathway (ΔG_direct_) or via the lipid membrane (ΔG_1_ + ΔG_2_). Values for protonated fentanyl (green) and protonated morphine (orange) are taken from the PMF calculations in Fig 1D and 4. Diffusion through the solvent (ΔG_3_) is assumed to be 0. **B.** Comparison of the free energy of binding to MOPr directly via the aqueous solvent, or indirectly via the membrane, where ΔG_indirect_ = ΔG_3_ + ΔG_2_ + ΔG_3_. **C.** 2D representation of the indirect, lipid binding route, using the same values as Fig 5A. Fentanyl (green) has a greater propensity to move into the lipid from the solvent, than morphine (orange).

As can be seen in Fig 5B, this indeed holds up, and the energies we have obtained here agree whether measured for the direct binding route or the indirect route, via the membrane. Importantly, whilst the overall binding energy for each ligand is very similar, the primary difference is the increased preference of fentanyl to partition into the lipid membrane (Fig 5C) where it can access the lipophilic access route. This suggests that fentanyl may favour this indirect, lipid access route, whereas morphine, which does not penetrate into the lipid, favours the “canonical” pathway, binding directly from the aqueous solvent.

## Discussion

Here, we applied CG MD simulations to study the interactions of both fentanyl and morphine with the MOPr. Using a combination of unbiased MD simulations and free energy calculations, we demonstrate that fentanyl exhibits a marked preference to access the MOPr orthosteric site via a novel binding route through the lipid membrane and MOPr transmembrane helices (Fig 6). Whereas, morphine accesses the orthosteric pocket by diffusing directly from the aqueous solvent and an extracellular vestibule site. Free energy calculations show that whilst fentanyl can also bind to the MOPr via the canonical aqueous route, fentanyl’s preference for the lipid access route is driven by its high lipid solubility; fentanyl rapidly partitions into the lipid membrane and clusters around the MOPr. In contrast, morphine only diffuses as far as the lipid surface. Once positioned at the TM6/TM7 lipid-facing interface, fentanyl interacts with hydrophobic and aromatic residues to induce formation of a gap through which it gains access to the MOPr orthosteric site.

**Figure. 6.**
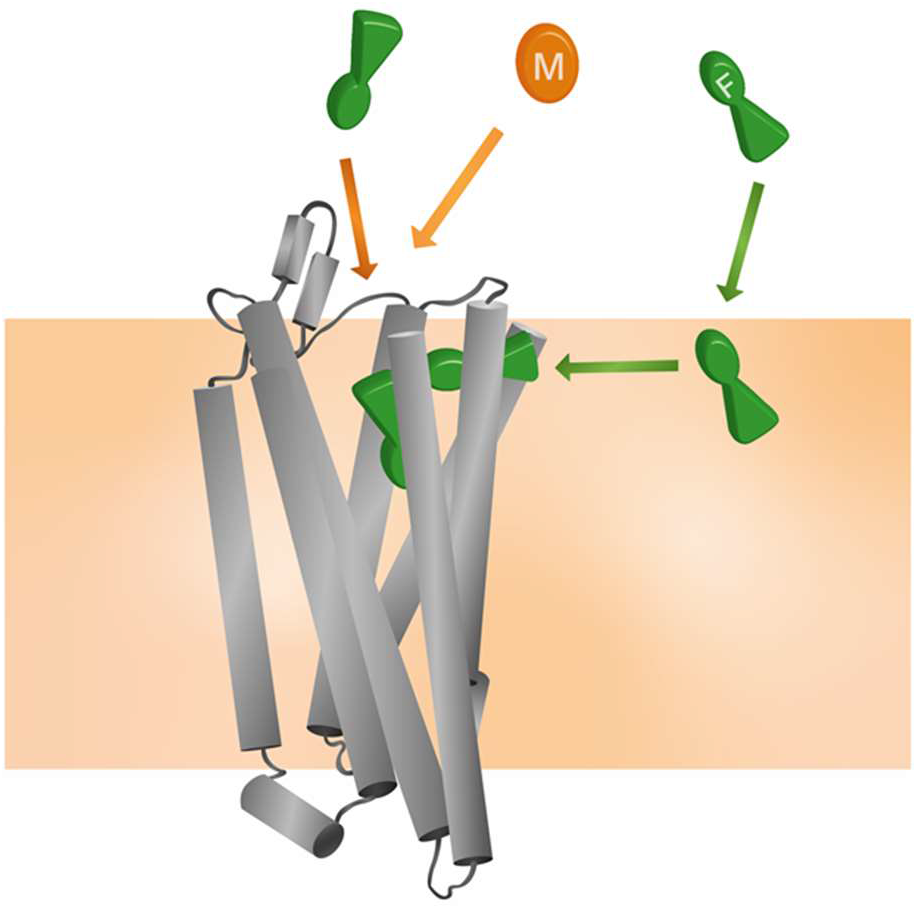
Model for the unique pharmacology of fentanyls at the MOPr. In competition with a morphinan ligand (such as morphine or naloxone), fentanyl (green) can access the orthosteric pocket via two binding routes; the canonical aqueous pathway and by the novel lipid pathway. In contrast, the morphinan ligand (orange) only has access to one binding route.

By using CG representation of our MOPr-ligand systems, we have captured ligand binding to the MOPr in a truly unbiased fashion, without the need for very longtime scale simulations (32) or use of any external potential or metadynamics approach (31). Combining our unbiased CG trajectories with steered MD and umbrella sampling to calculate the free energy landscapes of the binding events has proved a powerful tool for interrogating opioid ligand binding pathways.

Due to the lower resolution of the coarse-grained MD employed in this study, the two ligands represent multiple “fentanyl” or “morphinan” molecules. It is likely that other fentanyl analogues with high lipophilicity also exhibit lipid phase binding to the MOPr, for instance carfentanil, sufentanil and ohmefentanyl. The size of the fentanyl-induced gap between TM6 and 7 would suggest that fentanyl’s ability to bind via the lipid is a property of both its high lipophilicity and the elongated, flexible structure. Morphine, which is less lipid soluble, does not penetrate into the lipid far enough to access the gap, and is therefore unlikely to favour this binding pathway.

It is increasingly appreciated that GPCR ligands can bind to a variety of topographically distinct sites within the receptor structure (49). Allosteric binding pockets have been identified within the extracellular vestibule (50, 51), lipid-facing portions of the transmembrane helices (52), and the intracellular G protein coupling region (53), spanning the entire bilayer (54). Similarly, metastable sites populated as ligands bind and unbind between the orthosteric pocket and the aqueous phase have been identified, and it seems likely that for Class A GPCRs which recognise small molecule ligands, some of these sites may be conserved. Indeed, the extracellular vestibule to which morphine initially binds in our MD simulations appears to be analogous to the vestibule sites in the β1 and β2 adrenergic (32) and M3 muscarinic receptors (33). In extensive, all-atom simulations of MOPr in the presence of a high concentration of oliceridine, the ligand was also observed to bind to sites in the extracellular portion of the receptor (30).

A lipid phase binding route has been proposed for other GPCRs; notably rhodopsin and the CB2 cannabinoid, sphingosine-1-phosphate, PAR1 and P2Y1 receptors (34–37, 55), though not so far for the MOPr which has evolved to recognise non-lipophilic peptide ligands. Like fentanyl at MOPr, 2-arachidonoylglycerol and vorapaxar are reported to access the orthosteric pocket via the TM6/7 interfaces of the CB2 and PAR1 receptors, respectively (34, 35). Particularly, in simulations of vorapaxar unbinding from the PAR1 receptor, the ligand also exits via a gap formed by TM6/7 and ECL3, towards the extracellular side of the receptor (35). Similar to the MOPr, this gap is lined by small hydrophobic and polar residues and an aromatic residue in position 7.35 (tryptophan in MOPr, tyrosine in PAR1). In the CB2 receptor, the ligand entry gap is further towards the intracellular side of TM6 and 7 (34).

Interestingly, the novel binding mode we observe here may be specific to the MOPr over the δ-opioid receptor (DOPr). Fentanyl is highly selective for MOPr over the DOPr (56). Where the residues lining the TM6/7 gap in the MOPr are largely small polar, hydrophobic or aromatic side chains, in the DOPr ECL3 contains two positively charged and bulky arginine residues (R291 and R292) which may impede fentanyl binding by both repulsion of the protonated nitrogen and steric hinderance.

This novel mechanism of interaction with MOPr is of great importance to our understanding of the pharmacology of fentanyl.

Firstly, by concentrating fentanyl in the lipid membrane, the apparent concentration around the receptor is markedly increased, as the membrane acts as a reservoir. This high local concentration increases the likelihood of receptor association. Therefore, whilst morphine and fentanyl have very similar binding energies for MOPr, the actual likelihood of fentanyl binding would be far higher, and this might well explain the increased potency of fentanyl over morphine, particularly in cells where a complete, intact cell membrane is present.

Secondly, once fentanyl has partitioned into the bilayer it will switch from 3D diffusion in the solvent to 2D, lateral diffusion in the membrane (57). This reduction in dimensionality results in fentanyl having a greater chance of finding the receptor target, compared to morphinan ligands exhibiting 3D diffusion in the aqueous phase. Similarly, the membrane may also serve to organise the fentanyl molecules at a depth and orientation which favours MOPr binding through the TM6/7 interface (54).

Our identification of the TM6/7 metastable interface on the outside of the MOPr helices also invites the possibility that fentanyl exhibits “exosite” binding and re-binding, as described by Vauquelin & Charlton (58). Unlike morphinan ligands which bind and unbind via the aqueous phase, fentanyl is not free to diffuse away from MOPr and instead binds to the “exosite” TM6/7 interface. From here, fentanyl can then rapidly and efficiently rebind to the orthosteric site.

The mechanisms outlined here may also explain the poor reversibility of fentanyls by the morphinan antagonist naloxone. Naloxone has similar lipid solubility to morphine and is therefore unlikely to concentrate in the bilayer or have access to the lipid phase binding route and TM6/7 exosite (Fig 6). It would therefore only compete with fentanyl for binding via the aqueous route, not the lipophilic route. Whilst naloxone can still compete with fentanyl to occupy the orthosteric pocket, fentanyl can remain bound to the TM6/7 exosite and thus rapidly rebind to the orthosteric site once naloxone has dissociated. A similar phenomenon has been demonstrated for the lipophilic β2 adrenergic receptor agonist, salmeterol, where the ligand may be retained in the lipid membrane allowing reassertion of its agonist effects after wash-out (59, 60).

Fentanyl and other synthetic opioid agonists are driving the current opioid overdose epidemic in the United States (61). Fentanyl’s rapid onset and high potency are compounded by poor naloxone-reversibility, making the risk of fentanyl overdose high. Only by understanding fully how fentanyl interacts with and activates MOPr will we be able to develop better antagonists. We have previously shown that fentanyl-induced respiratory depression in mice is poorly reversed by naloxone compared to that by morphine. In light of these MD data, this is hardly surprising given that naloxone has low lipid solubility. However, we have also shown that the more lipophilic antagonist diprenorphine is better able to antagonize the effects of fentanyl (11). This might suggest that diprenorphine can at least access the entry point in the TM domains to block fentanyl access. Whilst elucidating how diprenorphine and other lipophilic ligands interact with the MOPr requires further study, the development of lipophilic MOPr antagonists may prove highly beneficial in combatting fentanyl overdose.

## Methods

### System set-up

The MOPr model was taken from the inactive, antagonist-bound crystal structure (26) (PDB: 4DKL), with the T4 lysozyme and ligands removed, and the missing intracellular loop 3 modelled using Insight II, as described in (45). The protein structure coordinates were then converted to coarse-grained MARTINI 2.2 representation using the *martinize* script (62). The secondary structure was constrained using an elastic network between backbone (BB) beads; elastic bonds with a force constant of 100 kJ mol^-1^nm^-2^ were defined between BB_i_-BB_i+4_ helix atoms, BBi-B B_i+10_ helix atoms, and BB atom pairs with low root mean square fluctuation and highly correlated motion as determined from all-atom MD simulations. All MD simulations were run using GROMACS 2019.2 (63).

To parameterise morphine and fentanyl in MARTINI, firstly, 1 μs all-atom MD simulations of fentanyl or morphine in water and 0.15 M NaCl were conducted under the Amber ff99SB-ildn force field (64). Ligands were parameterised using acpype/Antechamber and the General Amber Force Field (65). Atom-to-bead mapping for morphine and fentanyl was then created as shown in Supplementary Fig 1A and B, respectively. The CG ligands were then solvated in water and 0.15 M NaCl, energy minimized for 10000 steps using the steepest descents algorithm, box dimensions and temperature equilibrated, and then production MD was run for 1 μs. Bond lengths and angles were measured and compared to the all-atom simulations, to determine appropriate mapping and bonded terms.

### Unbiased CG simulations

The CG MOPr model was then embedded in a POPC:POPE:cholesterol lipid bilayer (ratio 5:5:1) using the *insane* script (66), and solvated in water, 0.15 M NaCl and 6 molecules of opioid ligand. The starting size of the system box was 15 x 15 x 15 nm^3^. Systems were first energy minimized over 50000 steps using the steepest descents algorithm, then equilibrated under NVT ensemble and then NPT ensembles, before production MD simulations were run at 310 K with a 10 fs timestep. The temperature and pressure were controlled by the V-rescale thermostat and Parrinello-Rahman barostat, respectively. Simulations were performed for up to 5 μs; the exact simulation lengths for each ligand are shown in Supplementary Table 1.

All simulations were analysed using the GROMACS suite of tools (63). Unless otherwise stated, all analyses were performed using the entire production trajectories. Data were plotted in GraphPad Prism v8, and images made in VMD (47).

### Free energy calculations

For the membrane/solvent partitioning calculations, systems were set up with small (5 x 5 x 10 nm^3^) membrane patches containing 32 POPE, 32 POPC and 6 cholesterol molecules, and solvated in 0.15 M NaCl. One molecule of either protonated fentanyl, neutral fentanyl, protonated morphine or neutral morphine was placed in the bilayer center. The systems were minimized for 50000 steps, keeping the ligand restrained. To generate the starting conformations for umbrella sampling, steered MD simulations were performed. Ligands were pulled from the bilayer center into the solvent (46), in a direction defined by the vector between the centers of mass of the ligand and the PO4 lipid beads, at a rate of 0.1 nm ns^-1^ and a force constant of 1000 kJ mol^-1^ nm^-2^.

For the ligand binding calculations, the final frames from the unbiased CG simulations with morphine or fentanyl bound in the orthosteric pocket were taken as the starting conformations. All other unbound ligands were removed, and the receptor-ligand complex was re-embedded in a smaller lipid bilayer (10 x 10 x 10 nm^3^). Steered MD simulations were performed to generate the starting conformations for umbrella sampling. In each case, separate simulations were performed to pull morphine or fentanyl from the orthosteric pocket along a) the aqueous / extracellular route, and b) the lipophilic / transmembrane domain route. The reaction coordinate was defined as the distance between the center of mass of the ligand and the receptor. Ligands were pulled at a rate of 0.1 nm ns^-1^ and a force constant of 1000 kJ mol^-1^ nm^-2^, with a 1000 kJ mol^-1^ nm^-2^ position restraint on 4 backbone beads (D114^2.50^, D147^3.32^, N150^3.35^ and S154^3.39^) of the MOPr to prevent translation or rotation of the receptor.

The starting conformations for umbrella sampling were extracted from these steered MD trajectories at 0.05 nm intervals along the reaction coordinate, generating ~80 umbrella sampling windows for each calculation. Each was subjected to 1 μs MD simulations, with a harmonic restraint of 1000 kJ mol^-1^ nm^-2^ to maintain the separation between the centers of mass of the ligand and PO4 beads (membrane partitioning calculations) or protein (ligand binding calculations). The PMFs were then extracted using the Weighted Histogram Analysis Method (WHAM) in GROMACS (67). PMFs were plotted as the average profile with statistical error calculated from bootstrap analysis. For the ligand binding calculations, ΔG_binding_ for each ligand in each binding pathway was calculated as the difference between the ligand-bound and final unbound states.

## Acknowledgements

The work described in this paper was supported by a grant from the Medical Research Council (MR/S010890/1) to GH, EK and SJC and was carried out using the computational facilities of the Advanced Computing Research Centre, University of Bristol http://www.bris.ac.uk/acrc/. We thank Roseanna Jackson of Slowe Club for artwork.

## Author contributions

KJS and RAC designed and performed the MD simulations. KJS, RAC and RBS analysed the data. GH, EK, SJC and KJS conceived the study. All authors contributed to and approved the content of the final manuscript.

## Supplementary Information

**Supplementary Table 1.**
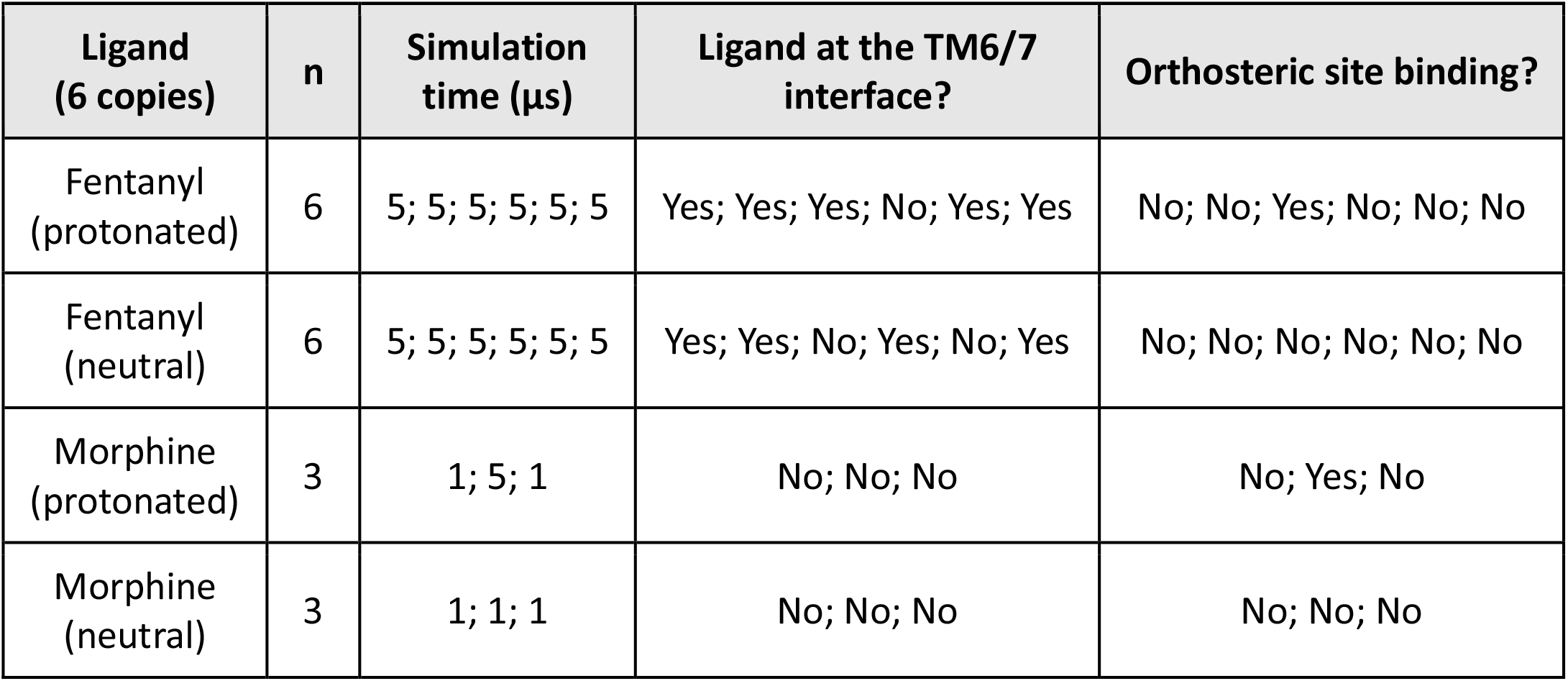
Long-timescale, independent CG simulations for each opioid ligand.

**Supplementary Figure 1.**
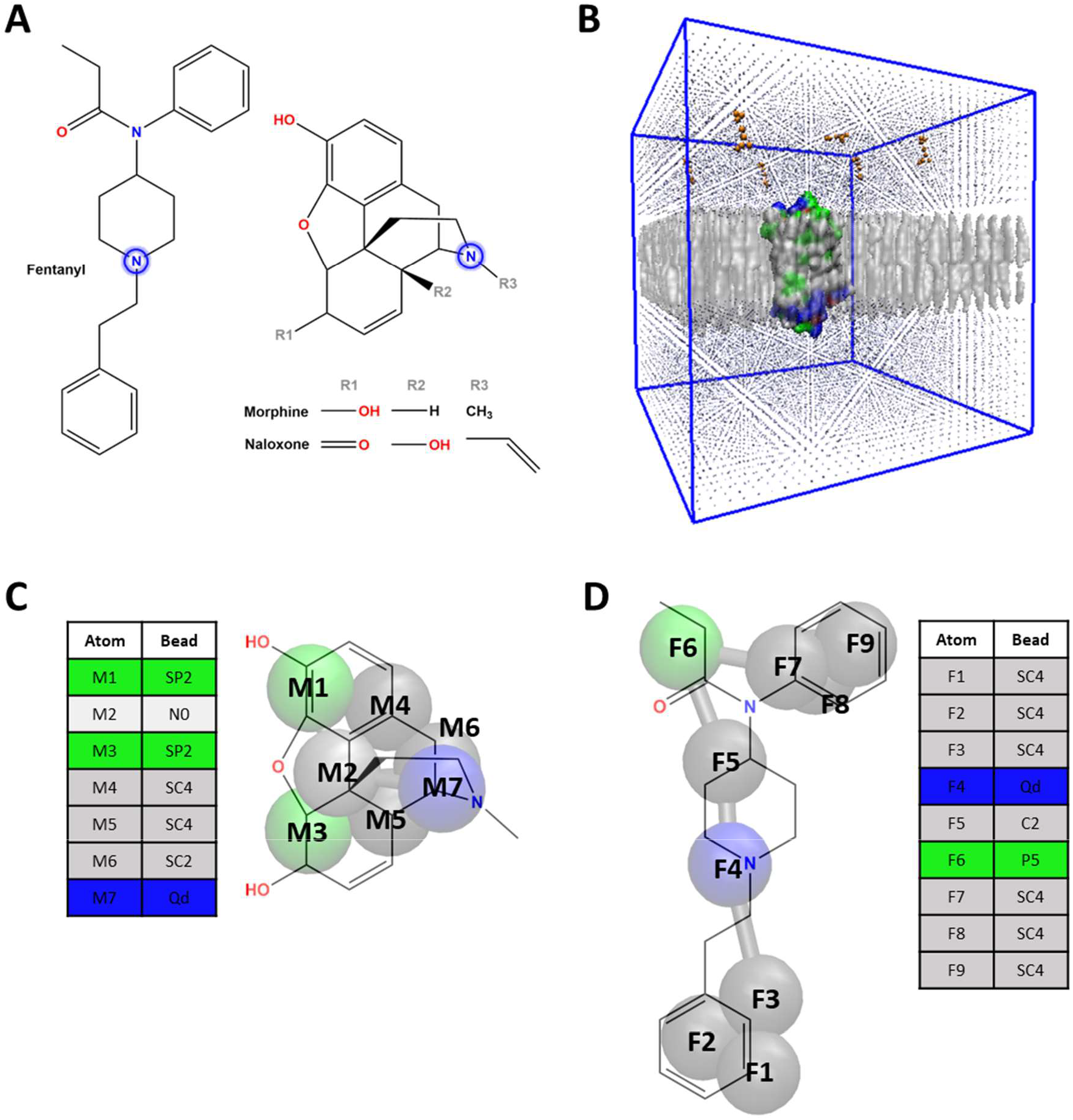
Ligand parameterization and system set up. **A.** Elongated structure of fentanyl compared to the rigid ring structures of morphine and naloxone. The protonatable amine in both molecules is highlighted. **B.** The systems were set up with CG MOPr embedded in a POPE, POPC, cholesterol membrane (grey), solvated in water and ion beads (blue) and 6 molecules of either fentanyl or morphine (orange) randomly placed in the solvent. **C.** Morphine was parameterized for the Martini forcefield using 7 Martini beads. **D.** Fentanyl was parameterized for the Martini forcefield using 9 Martini beads. For simulations of neutral (unprotonated) ligands the Qd beads were replaced with Nd beads. For explanation of MARTINI bead types, see *Marrink et al 2007*.

**Supplementary Figure 2.**
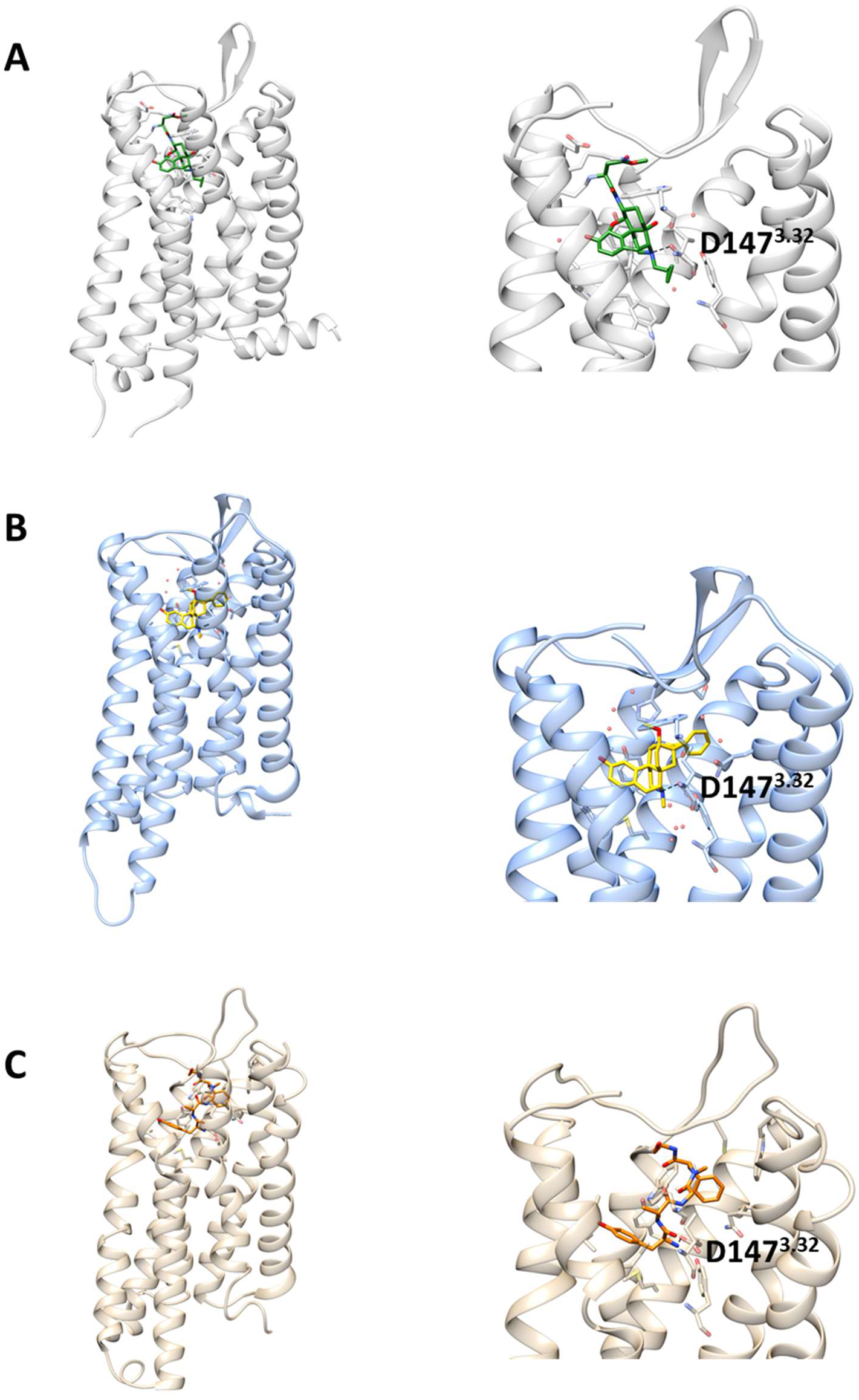
Structures of the MOPr. **A.** X-ray crystal structure of MOPr (grey) bound to the antagonist β-FNA (green). PDB 4DKL (*Manglik et al. 2012*). **B.** X-ray crystal structure of MOPr (blue) bound to the agonist BU72 (yellow). PDB 5C1M (*Huang et al. 2015*). **C.** Cryo-EM structure of MOPr (tan) bound to the peptide agonist DAMGO (orange). PDB 6DDF (*Koehl et al 2018*). In each case, residues forming the binding pocket are displayed as sticks and the key amine-D147^3.32^ interaction is indicated with a dashed line.

**Supplementary Figure 3.**
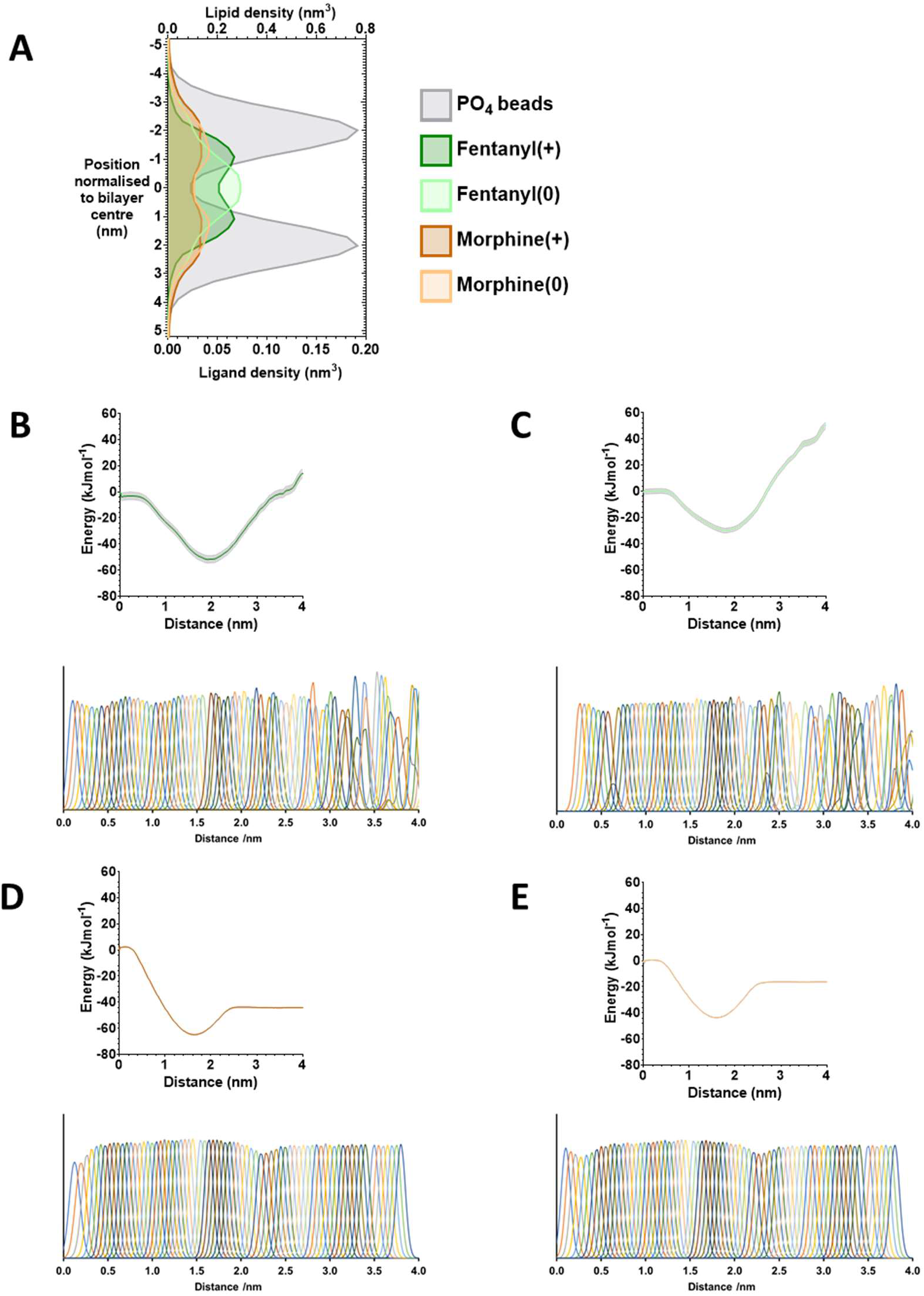
Free energy calculations for ligand solvent/membrane partitioning. **A.** Density plots showing the average position of protonated fentanyl (dark green), neutral fentanyl (light green), protonated morphine (dark orange) and neutral morphine (light orange), in relation to the phosphate beads of the lipid membrane (grey). **B-D.** Full PMF profiles along the reaction coordinate for the umbrella sampling simulations of ligands partitioning between the centre of the lipid bilayer (0 nm) and the bulk aqueous solvent (4 nm): **B.** protonated fentanyl, **C.** neutral fentanyl, **D.** protonated morphine and **E.** neutral morphine. The histograms alongside each PMF profile indicate the umbrella sampling windows fully captured the entire reaction coordinate.

**Supplementary Figure 4.**
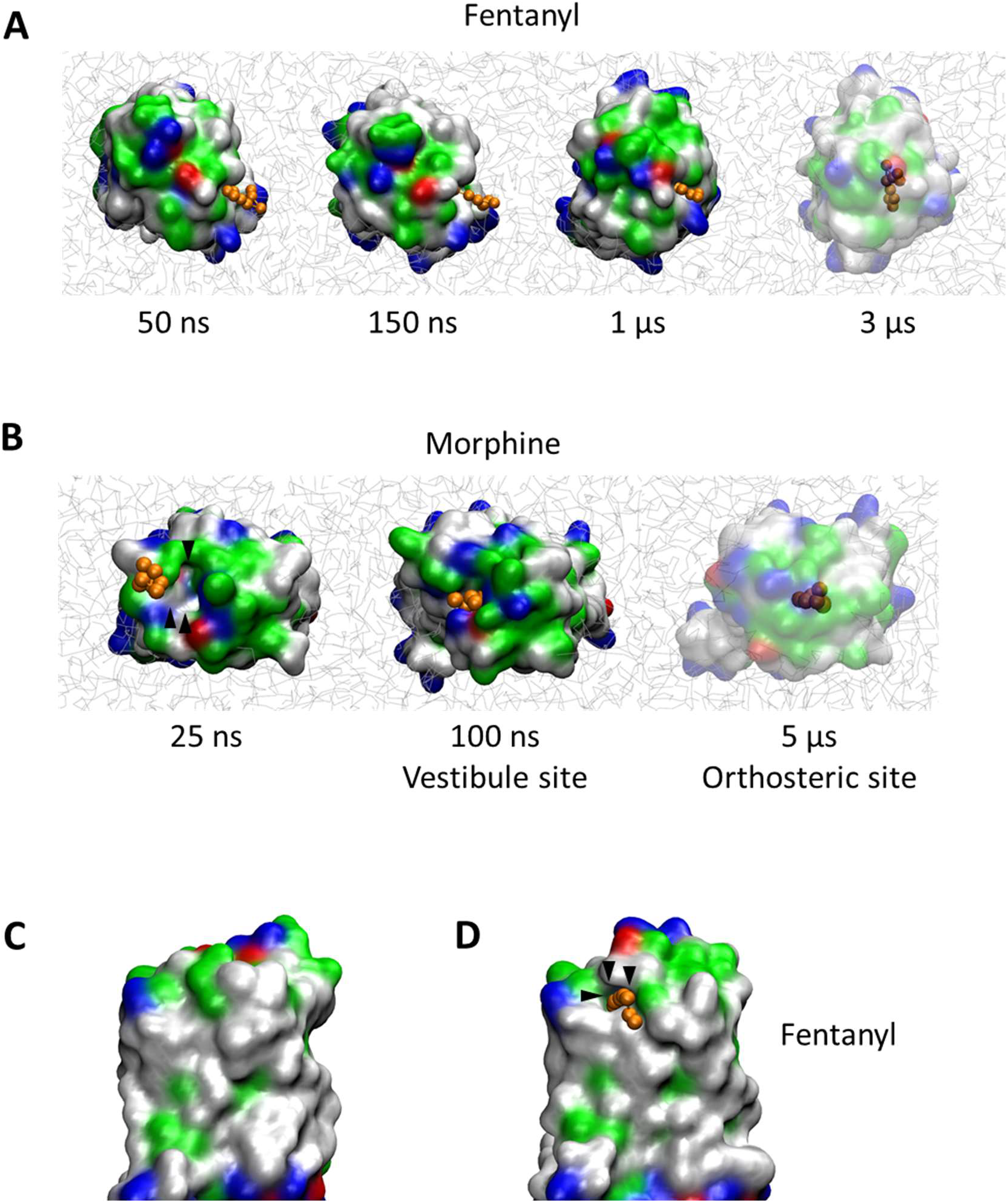
Opioid ligand binding to the MOPr. **A.** Fentanyl (orange) binding via the lipid membrane, with the MOPr viewed from the extracellular side of the membrane. **B.** Morphine (orange) binding via the extracellular vestibule (arrows), with MOPr viewed from the extracellular side of the membrane. **C.** Surface representation of the TM6/7 interface in the absence of fentanyl. **D.** Fentanyl induces formation of a gap between TM6/7 (arrows) through which the ligand can bind. The protein is coloured according to residue properties (hydrophobic; grey, polar; green, acidic; red, basic; blue).

**Supplementary Figure 5.**
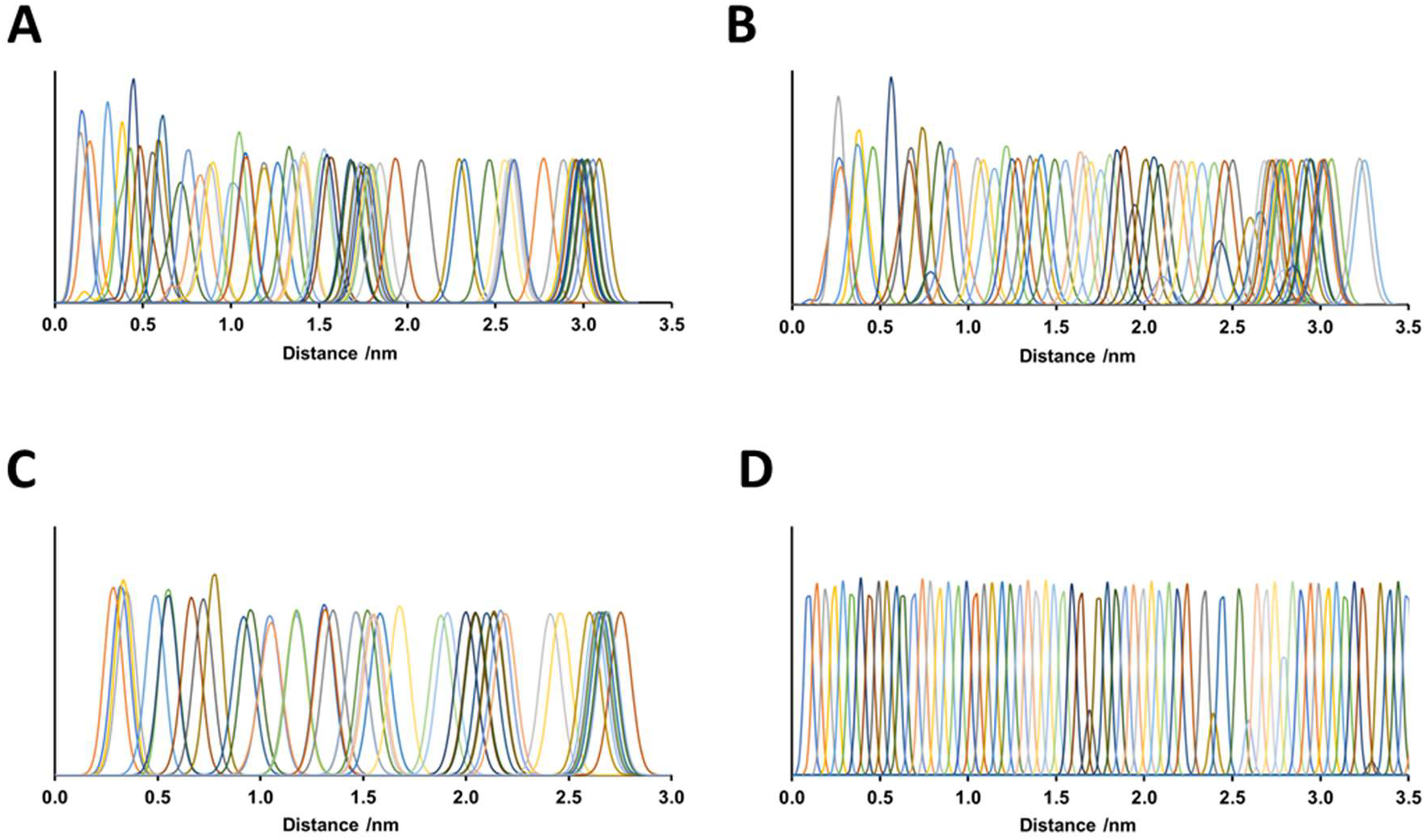
Free energy calculations for ligand binding pathways. **A.** Histograms along the reaction coordinate for the umbrella sampling simulations of morphine binding via the aqueous pathway. **B.** Histograms along the reaction coordinate for the umbrella sampling simulations of fentanyl binding via the aqueous pathway. **C.** Histograms along the reaction coordinate for the umbrella sampling simulations of morphine binding via the lipid pathway. **D.** Histograms along the reaction coordinate for the umbrella sampling simulations of fentanyl binding via the lipid pathway.

**Supplementary Movie 1. Fentanyl binding via the lipid bilayer and transmembrane domains**

**Supplementary Movie 2. Morphine binding via the aqueous solvent and extracellular vestibule**

